# Room without a view – den construction in relation to body size in brown bears

**DOI:** 10.1101/865188

**Authors:** Shotaro Shiratsuru, Andrea Friebe, Jon E. Swenson, Andreas Zedrosser

## Abstract

Hibernation is an adaptive strategy to survive harsh winter conditions and food shortage. The use of well-insulated winter dens helps animals minimize energy loss during hibernation. Brown bears (*Ursus arctos*) commonly use excavated dens for hibernation. Physical properties of excavated dens, such as the amount of space between a bear and the inner wall, wall/roof thickness, and bedding materials, are expected to impact heat retention and energy conservation of bears. The objective of this study was to examine the impact of physical properties of excavated dens on energy conservation in hibernating bears. Our hypothesis was that bears excavate dens in a way to minimize heat loss and optimize energy conservation during hibernation. We predicted that physical properties of excavated dens would significantly affect the bears’ post-hibernation body condition. To test our hypothesis and prediction, we analyzed data collected from brown bears in Sweden with linear mixed effects models, examining (i) what factors affect den-excavation behavior and (ii) if physical properties of excavated dens affect post-hibernation body condition. We found that bears excavated a den cavity in relation to their body size, that older bears tended to excavate better-fitting den cavities compared to young bears, and that the physical properties of excavated dens did not significantly affect a bears’ post-hibernation body condition. Older bears excavated better-fitting den cavities, suggesting a potentially experience-based shift with age in den-excavation behavior and an optimum cavity size relative to a bear’s body size. The strong year effect shown by the most parsimonious model for post-hibernation body condition suggests that variations in physical properties of excavated dens are possibly negligible, compared to the large annual variations in biotic and abiotic factors affecting pre-hibernation body condition and heat loss during hibernation.

## Introduction

Hibernation is a physiological and behavioral adaptation through which animals survive harsh seasonal conditions, such as inclement weather or low food availability, by minimizing energy loss [1-3]. Small mammalian hibernators, such as arctic ground squirrels (*Spermophilus parryii*) and Alpine marmots (*Marmota marmota*), decrease their body temperatures to around 0 C or even lower during hibernation to overcome their high mass-specific metabolic rates and low amount of body fat stores. On the other hand, large mammalian hibernators with large amount of body fat stores, such as brown bears (*Ursus arctos*) and American black bears (*U. americanus*), decrease metabolic rates while maintaining relatively high body temperatures [4-5].

Most brown bears and American black bears spend 4-6 months in winter dens without eating or drinking [3, 6, 7], while using the fat storage gained during hyperphagia as their main energy source and conserving lean body mass via urea recycling [8-13]. In addition, bears give birth during hibernation and the cubs are fed on milk produced from the stored fat and lean body mass [12, 14, 15]. To cope with this exclusive dependence on stored fat and protein reserves for survival and reproduction during hibernation, in addition to the slow cooling rate of the body due to their relatively small surface area to volume ratio [4], bears use metabolic inhibition independently of body temperature [13, 16, 17]. American black bears suppress their metabolism to 25% of the summer basal metabolic rate, but body temperature only decreases from 37-38 C to an average of 33 C in mid-hibernation [13]. This mechanism enables bears to reduce the thermal gradient between the body and the environment, thereby minimizing energy loss [18]. Hibernating bears shiver to produce extra heat in cold ambient and den temperatures, thereby inducing cycles of body temperatures [17]. Therefore, the use of well-insulated dens should help bears minimize energy loss during hibernation and bears should select dens optimally in relation to energy conservation [19].

The amount of protection and insulation provided by a den may vary depending on the den type and differences potentially influence the amount of heat loss and vulnerability to disturbances, thereby potentially affecting the bears’ survival and reproduction [20, 21, 22]. Enclosed dens, such as tree or rock cavities and excavated dens, offer protection and insulation from inclement weather [1, 2, 20, 23] and thus are likely to be preferred by bears. Especially in excavated dens, which can be adjusted by an individual in relation to its body size, radiant heat from the soil and metabolic heat from the bear can be trapped within the den and keep the den temperature higher than the ambient temperature [6, 24, 25]. Bedding materials on the ground may enhance insulation, by forming a microclimate between the bear and the soil [26-28]. Consequently, enclosed dens provide bears with a microenvironment where temperatures are relatively warm and stable, compared to outside temperatures, thereby optimizing energy conservation [27, 28]. The tendency of female bears to select for enclosed dens [2, 3, 29] can be explained by the high energy demand of females for birth and lactation during the denning period [12]. Female bears utilizing excavated dens have higher reproductive success compared to those using other den types [28, 30]. However, to our knowledge, there are no studies evaluating the potential impact of the physical properties of enclosed dens, such as the size of the den cavity in relation to a bear’s body size, wall thickness, and bedding materials, on energy loss in hibernating bears.

Worldwide, brown bears mainly use excavated dens for hibernation [21, 27]. Den cavity size, composition of the wall/roof (from now on referred to as den composition), wall/roof thickness, and bedding materials have been proposed as important factors that influence heat retention and energy conservation, thereby determining the quality of an excavated den [1, 25, 31]. A bear’s body size has been suggested to determine den cavity size [23, 31]. The volume of the air space between a bear and the cavity wall likely varies, with greater air space within the den resulting in increased convective heat loss caused by enhanced air flow [1, 28, 31]. However, an optimum size of an air space warmed by the bear’s radiative heat could contribute to efficient heat retention [21, 31]. Wall/roof thickness may be important for preserving heat within the den [25, 29].

The objective of this study was to examine how energy conservation in bears is affected by the physical properties of excavated dens, based on the hypothesis that bears excavate dens to minimize heat loss and optimize energy conservation during hibernation. First, we explored what factors affect den construction behavior in bears, focusing on the size of the den cavity and the volume of the air space between a bear and the cavity wall. We predicted that bears would excavate den cavities in relation to their body size. We also predicted that neither sex nor age of bears would affect the volume of the air space between their bodies and the cavity wall, assuming that bears try to minimize the air space in the den cavity to prevent convective heat loss. We then examined if physical properties of excavated dens affect energy conservation during hibernation. We predicted that energy conservation in hibernating bears is positively related to wall thickness and size of the bedding materials, but negatively related to the volume of the air space between a bear and the cavity wall in relation to a bear’s body size.

## Materials and methods

### Study area

The study area was in Dalarna and Gävleborg counties in south-central Sweden (∼13,000 km^2^, ∼61N, 14E). The rolling terrain is covered by an intensively managed forest and elevation ranges from 200 m in the southeast to 1,000 m in the west. Average temperature is -7 C in January and 15 C in July, and snow cover generally lasts from late October until early May. The mean annual precipitation is 600-1,000 mm, and the vegetation period ranges from 150-180 days [33]. The area is mainly covered by Scots pine (*Pinus sylvestris*) and Norway spruce (*Picea abies*) interspersed with deciduous trees, such as mountain birch (*Betula pubescens*), silver birch (*B. pendula*), aspen (*Populus tremula*), and gray alder (*Alnus incana*). Ground vegetation consists of mosses, lichens, grass, heather and berries, including bilberries (*Vaccinium myrtillus*), lingonberries (*V. vitis-idaea*), and crowberries (*Empetrum hermaphroditum*), which are the main foods of bears in autumn [34].

Brown bears in Scandinavia hibernate in dens from late October to late April, although males spend less time in dens than females, and the denning duration varies in relation to age and reproductive status in females [7, 22]. In central Scandinavia, ants from the family *Formica* build very large mound-shaped nests, and abandoned “anthills” overgrown by berry bushes, can be excavated and used by bears as winter dens. Anthill dens are the most common winter dens among brown bears in central Scandinavia, utilized by 56% of females and 54% of males [22]. This high use of anthill dens can be explained by the high abundance and the high insulating effect of anthills, and females hibernating in anthill dens tend to have a higher reproductive success [22, 30, 35]. “Soil dens” are the dens excavated in soil [22]. So-called “nest dens”, where bears only collect a protective layer of bedding material on the ground, but are otherwise exposed to the elements, are most commonly used by adult males [36].

### Data collection and preparation

Bears were immobilized by darting from a helicopter in spring shortly after den exit and fitted with VHF (Very High Frequency) radio transmitters (1985-2002) or GPS (Global Positioning System) - GSM (Global System for Mobile Communication) collars (2003-present) [30, 37] by the Scandinavian Brown Bear Research Project (SBBRP, www.bearproject.info). Bears were not captured before den entry to avoid potential disturbance, according to accepted veterinary and ethical procedures. See Zedrosser et al. (2006) [37] and Arnemo et al. (2012) [38] for more detailed information on capture and handling.

Body length (cm) was measured with a tape measure as the length from the tip of the nose to the base of the tail, and chest circumference (cm) was measured at the widest part of the chest [37]. Body mass was measured to the nearest kg with a spring scale. Ages of bears that were not first captured as yearlings with their mothers were estimated by extracting a premolar tooth and counting cementum annuli [39]. Bears captured after 5 May were excluded from the analysis to avoid changes in weight or body condition after leaving the den, which might affect the results [40].

The SBBRP has collected data on winter dens from 1986 to 2016. Winter dens were categorized into 3 types, anthill dens, anthill/soil dens (20-80 % of the den material consisted of an anthill and the rest of soil), and soil dens (> 80% of the den material was soil) [41]. For each den, we recorded size (length x width x height) of the whole den (i.e., on the outside), as well as the size of the den cavity, wall/roof thickness, size of bedding materials (length x depth), and habitat information, such as the number of trees (>10 cm circumference at chest height) within a 10-m radius around a den. In this study, we only used data from solitary bears that used anthill, anthill/soil, and soil dens, and did not change dens during the winter.

Many bears have been captured and recorded multiple times in different years during our study. For the analyses of den cavity size, we used the data from 97 observations of 69 solitary bears. We used the data from 96 observations of 68 solitary bears for the analysis of the volume of the air space between a bear and the cavity wall in relation to a bear’s body size. For the analysis of post-hibernation body condition index, we used the data from 67 observations of 53 solitary bears.

### Data analysis

Whole den size (from now on referred to as den size) and cavity size were estimated based on the assumption that both the den and the cavity had the shape/volume of a half-dome. In addition, we calculated indices for the average thickness of the den wall and the size of the bed inside the den. Equations for each variable are as follows:

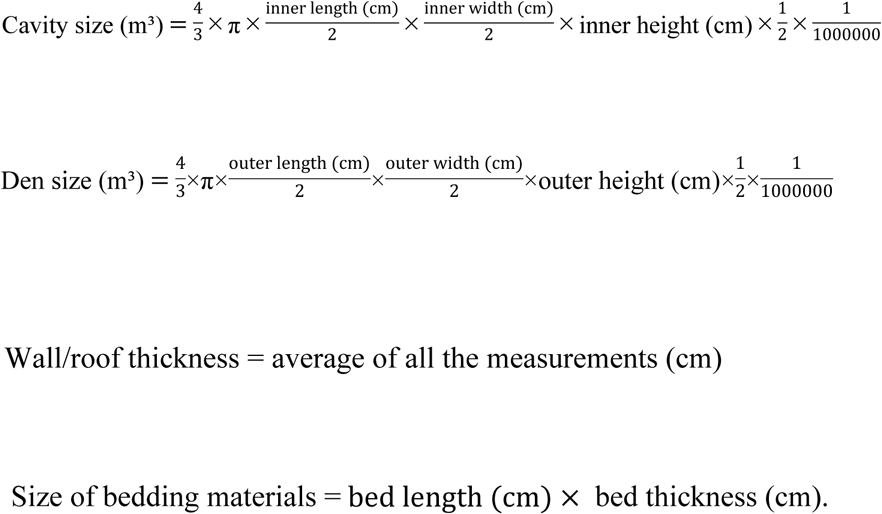

As an index of the volume of the air space between a bear and the cavity wall in relation to a bear’s body size, we calculated the ratio of body size to cavity size (body-cavity ratio) by estimating a bear’s body volume on the assumption that it resembles a cylinder. Equations used for calculating the body-cavity ratio are as follows:

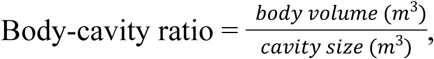

where

body volume 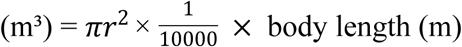, and

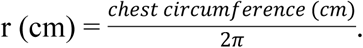

Because we only used individuals that were captured after hibernation in this analysis, loss of fat or body mass during hibernation could not be obtained. Instead, we used a post-hibernation body condition index (BCI) to evaluate the relative energy status of bears after hibernation [41], which can be considered as an index of energy conservation during hibernation. The BCI defines body condition as total body mass (kg) relative to body size (cm) [41]. We calculated BCI as the standardized residual from the linear regression of body mass (kg) against linear body length (cm). Both body mass and linear body length were log-transformed [41]. We confirmed that there is no correlation between the calculated BCI and linear body length (*r* = 0.031, *p* = 0.801, n = 68).

To test out hypothesis and predictions, we used linear mixed effects models to examine the impact of potential variables on 1) cavity size, 2) body-cavity ratio, and 3) post-hibernation BCI. We constructed a candidate model for cavity size by including age, sex, and body length as predictor variables. For body-cavity ratio, we included sex and age as predictor variables in a candidate model. In both analyses, we compared the candidate model with a null model, which did not include any predictor variables, to test if the candidate model was more parsimonious than the null model. Because some bears were sampled multiple times, we added individual ID into all the models as a random effect. In the analysis of post-hibernation BCI, several biotic and abiotic factors needed to be considered in addition to the physical properties of winter dens. Post-hibernation body condition is expected to be positively related to pre-hibernation body condition, which has been reported to increase with age [9, 11]. In addition, energetic costs and weight loss in bears generally increase with the duration of hibernation [15], and the duration of denning varies depending on sex and reproductive status [7, 22, 42]. Heat loss during hibernation can be exacerbated by severe winter temperatures [17], even if the animal is hibernating in an enclosed cavity [1]. In addition, the loss of energy and body mass of bears during hibernation is highly affected by pre-hibernation body condition [9, 15, 37]. Bears in Scandinavia rely mostly on berries, especially bilberries, for gaining fat reserves in autumn [33, 43, 44], therefore berry production has an impact on pre-hibernation body condition. Some studies have suggested the importance of snow deposition for insulation [23, 24, 45, 46]. We constructed two candidate models in the analysis of post-hibernation BCI. One of them included den composition, wall thickness, and the size of bedding materials as predictor variables, based on our hypothesis that physical properties of excavated dens affect heat loss of bears during hibernation. The other candidate model included only sex and age as predictor variables, based on an alternative hypothesis that physical properties of excavated dens would not have significant effects on energy conservation of hibernating bears. We compared these two candidate models with a null model that did not include any predictor variables. To control for the biotic and abiotic factors, which are highly variable from year to year, we included year as a random effect in addition to individual ID in all the models for post-hibernation BCI. Cavity size, body-cavity ratio, wall thickness, and size of bedding materials were log-transformed in all analyses to achieve homogeneity of variance and normal distribution of residuals [47, 48]. The software R 3.4.2 [49] was used for all analyses. In all the statistical analyses, the most parsimonious model was selected using Akaike’s Information Criterion corrected for small sample size (AICc) [50, 51] to obtain parameter estimates. Linear mixed effects models were analyzed with the *lmer* function in the *lmerTest* package [52], and model comparison was conducted with *AICcmodavg* package [53]. In each analysis, we excluded variables showing a variance inflation factor (VIF) greater than 3 [48]. We identified and removed outliers in predictor and response variables in each analysis by visualizing data with boxplots and the Cleveland dotplots [48].

## Results

The most parsimonious model explaining den cavity size included a bear’s body length, sex, and age as predictor variables (Table 1). Den cavity size increased significantly with a bear’s body length, but sex and age did not have significant effects on den cavity size (Table 1).

**Table 1.** Comparison of candidate linear mixed effects models predicting the size of den cavity (log transformed) in an excavated den of brown bears in Sweden during 1986-2016 (n=98 from 69 solitary bears) and parameter estimates from the most parsimonious model.

| Model comparison |  |  |  |  |  |
| --- | --- | --- | --- | --- | --- |
| Model | k | $\Delta AICc$ | | $w$ | |
| Body length + Sex + Age | 6 | 0 |  | 1 |  |
| ~1 | 3 | 36.85 |  | 0 |  |
| Parameter estimates from the most parsimonious model |  |  |  |  |  |
| Variable | $\beta$ | SE | Confidence interval | t | $p$ |
| Body length (m) | 1.94 | 0.33 | (1.28, 2.57) | 5.93 | < 0.001 |
| Sex: male | 0.14 | 0.12 | (-0.09, 0.36) | 1.16 | 0.25 |
| Age | -0.03 | 0.02 | (-0.06, 0.01) | -1.63 | 0.11 |

Both models included individual ID as a random effect. k is the number of parameters including intercept, $\Delta AICc$ is the change in AICc from the most parsimonious model, and w is
Akaike model weight. $\beta$ is the parameter estimate, SE is the standard error, confidence interval is 95% confidence interval, $t$ is the $t$ -value, and $p$ is the $p$ -value.

The most parsimonious model explaining body-cavity ratio included age and sex as predictor variables (Table 2). Body-cavity ratio increased significantly with age, but sex did not have a significant effect on body-cavity ratio (Table 2).

**Table 2.**
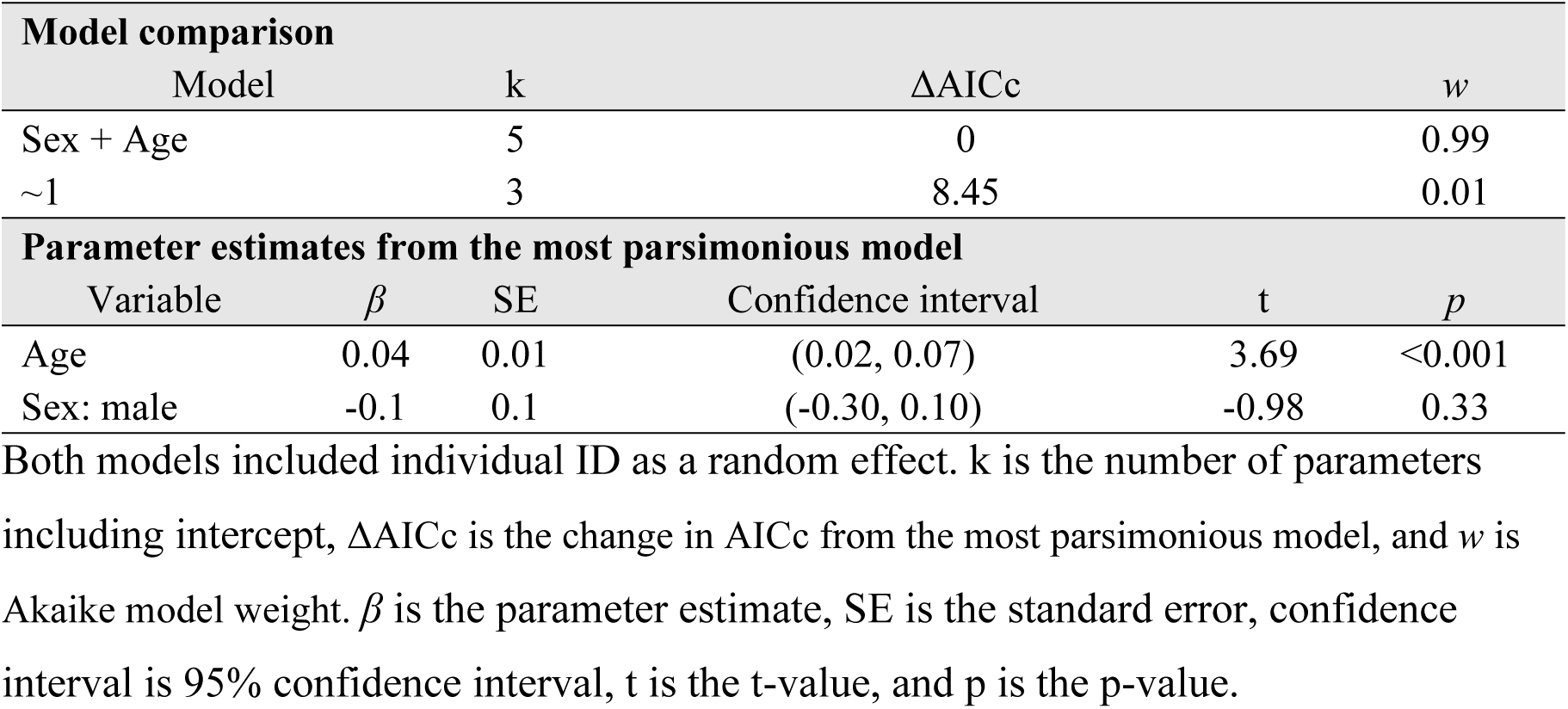
Comparison of candidate linear mixed effects models predicting body-cavity ratio (log transformed) of an excavated den of brown bears in Sweden during 1986-2016 (n=97 from 68 solitary bears) and parameter estimates from the most parsimonious model.

| Model comparison |  |  |  |  |  |
| --- | --- | --- | --- | --- | --- |
| Model | | k | $\Delta AICc$ | $w$ | |
| Sex + Age |  | 5 | 0 | 0.99 |  |
| ~1 |  | 3 | 8.45 | 0.01 |  |
| Parameter estimates from the most parsimonious model |  |  |  |  |  |
| Variable | $\beta$ | SE | Confidence interval | t | $p$ |
| Age | 0.04 | 0.01 | (0.02, 0.07) | 3.69 | <0.001 |
| Sex: male | -0.1 | 0.1 | (-0.30, 0.10) | -0.98 | 0.33 |

Both models included individual ID as a random effect. $k$ is the number of parameters including intercept, $\Delta AICc$ is the change in $AICc$ from the most parsimonious model, and $w$ is Akaike model weight. $\beta$ is the parameter estimate, SE is the standard error, confidence interval is 95% confidence interval, $t$ is the $t$ -value, and $p$ is the $p$ -value.

The most parsimonious model explaining post-hibernation BCI included age and sex as predictor variables (Table 3). Post-hibernation BCI increased significantly with age, but sex did not significantly affect post-hibernation BCI (Table 3). Significant interannual variations were indicated by the relatively large variance of year as a random effect (0.39). However, the large residual variance (0.47) suggests that a large portion of the variation in post-hibernation BCI could not be explained by the interannual variations and the predictor variables (sex and age).

**Table 3.** Comparison of candidate linear mixed effects models predicting the post-hibernation body condition index of brown bears in Sweden during 1986-2016 (n=67 from 53 solitary bears) and parameter estimates from the most parsimonious model.

| Model comparison |  |  |  |  |  |
| --- | --- | --- | --- | --- | --- |
| Model | | k | $\Delta AICc$ | $w$ | |
| Sex + Age |  | 6 | 0 | 0.92 |  |
| ~1 |  | 4 | 5.23 | 0.07 |  |
| Bed size + Wall thickness<br>+ Den type + Body-cavity<br>ratio |  | 9 | 7.95 | 0.02 |  |
| Parameter estimates from the most parsimonious model |  |  |  |  |  |
| Variable | $\beta$ | SE | Confidence interval | t | $p$ |
| Age | 0.05 | 0.02 | (0.01, 0.09) | 2.47 | 0.02 |
| Sex: male | 0.23 | 0.19 | (-0.14, 0.63) | 1.21 | 0.23 |

All models included individual ID and year as random effects. Bed size, wall thickness, and body-cavity ratio were log-transformed. k is the number of parameters including intercept, $\Delta AICc$ is the change in AICc from the most parsimonious model, and $w$ is Akaike model weight. $\beta$ is the parameter estimate, SE is the standard error, confidence interval is 95% confidence interval, t is the t-value, and p is the p-value.

## Discussion

Our main findings were that bears excavated a den cavity in relation to their body size, that older bears excavated better-fitting den cavities by reducing the amount of space between den wall and their bodies, and that physical properties of excavated winter dens did not have significant effects on the post-hibernation body condition of bears.

Den cavity size was positively related to a bear’s body size, as reported in previous studies [23, 28, 32]. Contradicting our prediction, the candidate model including age and sex as predictor variables was selected over the null model. The body-cavity ratio increased with age, implying that older bears excavated better-fitting den cavities. A potential explanation for this age effect is that older bears may be more experienced and skilled, and therefore able to excavate cavities that better fit their bodies to reduce heat loss during hibernation compared to younger and less experienced bears. Experience likely is an important factor that affects the behavior in bears. For example, seal hunting behavior of polar bears (*Ursus maritimus*) was reported to improve with age [54]. It has also been reported that older brown bears have higher yearly reproductive success than younger bears, probably because older bears with more experience are more competitive in mating and better at rearing offspring [55, 56].

Despite the use of a potential strategy to reduce heat loss during hibernation by bears, the physical properties of excavated dens did not affect post-hibernation BCI. The candidate model including volume of the air space between a bear and the cavity wall (body-cavity ratio), wall/roof thickness, den composition, and the size of bedding materials as predictor variables was not selected as the most parsimonious model (Table 3). Wall thickness and the air space within the den have been reported to be important for insulation and heat retention for denning animals [21, 25, 29, 31]. In addition, bedding materials on the ground have been suggested to enhance insulation [26-28]. However, we found no such effects. Instead, age showed a significant effect on post-hibernation BCI, in accordance with previous studies reporting that a bear’s pre-denning body condition is expected to increase with increasing age and thereby affects post-hibernation body condition [9, 11, 15]. The large interannual variance of post-hibernation BCI shown by the most parsimonious model suggests potential impacts of interannual variations in biotic and abiotic factors on post-hibernation body condition of bears. For example, severe winter temperatures [1, 17], berry production [33, 43, 44], and snow deposition [23, 24, 45, 46] are expected to affect pre-hibernation body condition and energy loss during hibernation [9, 15, 37], thereby affecting post-hibernation body condition. Sex may affect the energy loss during hibernation, likely because females spend significantly more time in dens than males [22], and energy cost and weight loss increase with longer duration of hibernation [15]. However, we did not find a relationship between sex and post-hibernation BCI. This is probably because the female bears in our data were all solitary. In general, solitary females do not hibernate as long as pregnant females [7, 22].

For future studies, it would be important to compare pre- and post-hibernation body conditions (fat and lean mass) of bears and take the length of denning into account to evaluate the actual influence of the properties of excavated dens on energy loss of denning bears. To do this correctly, it would be necessary to capture and measure bears before and after hibernation and to track the dates of den entry and exit. Moreover, the benefits of denning in excavated dens compared to fixed dens (e.g., rock cavity dens) in energy conservation in bears could be examined by comparing the amount of fat and lean mass loss between bears using excavated dens and those using other den types.

## Acknowledgements

We are grateful to the many field works and volunteers that have contributed to the data collection for this study during the course of the years. We especially acknowledge the help of Dr. honoris causa S. Brunberg. The long-term funding of the Scandinavian Brown Bear Research Project (SBBRP) has come primarily from the Swedish Environmental Protection Agency, the Norwegian Environment Agency, the Austrian Science Fund, and the Swedish Association for Hunting and Wildlife Management. This is paper No. XXX from the SBBRP.

